# The human genetic variant rs6190 unveils Foxc1 and Arid5a as novel pro-metabolic targets of the glucocorticoid receptor in muscle

**DOI:** 10.1101/2024.03.28.586997

**Authors:** Ashok Daniel Prabakaran, Hyun-Jy Chung, Kevin McFarland, Thirupugal Govindarajan, Fadoua El Abdellaoui Soussi, Hima Bindu Durumutla, Chiara Villa, Kevin Piczer, Hannah Latimer, Cole Werbrich, Olukunle Akinborewa, Robert Horning, Mattia Quattrocelli

**Affiliations:** Molecular Cardiovascular Biology, Heart Institute, Cincinnati Children’s Hospital Medical Center and Dept. Pediatrics, University of Cincinnati College of Medicine, Cincinnati, OH, USA; Stem Cell Laboratory, Department of Pathophysiology and Transplantation, Dino Ferrari Centre, University of Milan, Italy; Systems Biology and Physiology Graduate Program, University of Cincinnati College of Medicine, Cincinnati, OH 45229, USA

**Keywords:** Glucocorticoid receptor, rs6190, muscle metabolism, insulin sensitivity, glucose tolerance, fatty acid uptake

## Abstract

The genetic determinants of the glucocorticoid receptor (GR) metabolic action remain largely unelucidated. This is a compelling gap in knowledge for the GR single nucleotide polymorphism (SNP) rs6190 (p.R23K), which has been associated in humans with enhanced metabolic health but whose mechanism of action remains completely unknown. We generated transgenic knock-in mice genocopying this polymorphism to elucidate how the mutant GR impacts metabolism. Compared to non-mutant littermates, mutant mice showed increased insulin sensitivity on regular chow and high-fat diet, blunting the diet-induced adverse effects on adiposity and exercise intolerance. Overlay of RNA-seq and ChIP-seq profiling in skeletal muscle revealed increased transactivation of *Foxc1* and *Arid5A* genes by the mutant GR. Using myotropic adeno-associated viruses for in vivo overexpression or knockdown in muscle, we found that *Foxc1* was required and sufficient for normal expression levels of insulin response pathway genes *Insr* and *Irs1*, promoting muscle insulin sensitivity. In parallel, *Arid5a* was required and sufficient to transcriptionally repress the lipid uptake genes *C 36* and *Fabp4*, reducing muscle triacylglycerol accumulation. Moreover, the Foxc1 and Arid5a programs in muscle were divergently changed by glucocorticoid regimens with opposite metabolic outcomes in muscle. Finally, we found a direct human relevance for our mechanism of SNP action in the UK Biobank and All of Us datasets, where the rs6190 SNP correlated with pro-metabolic changes in BMI, lean mass, strength and glucose control according to zygosity. Collectively, our study leveraged a human nuclear receptor coding variant to unveil novel epigenetic regulators of muscle metabolism.

**Graphical abstract:** 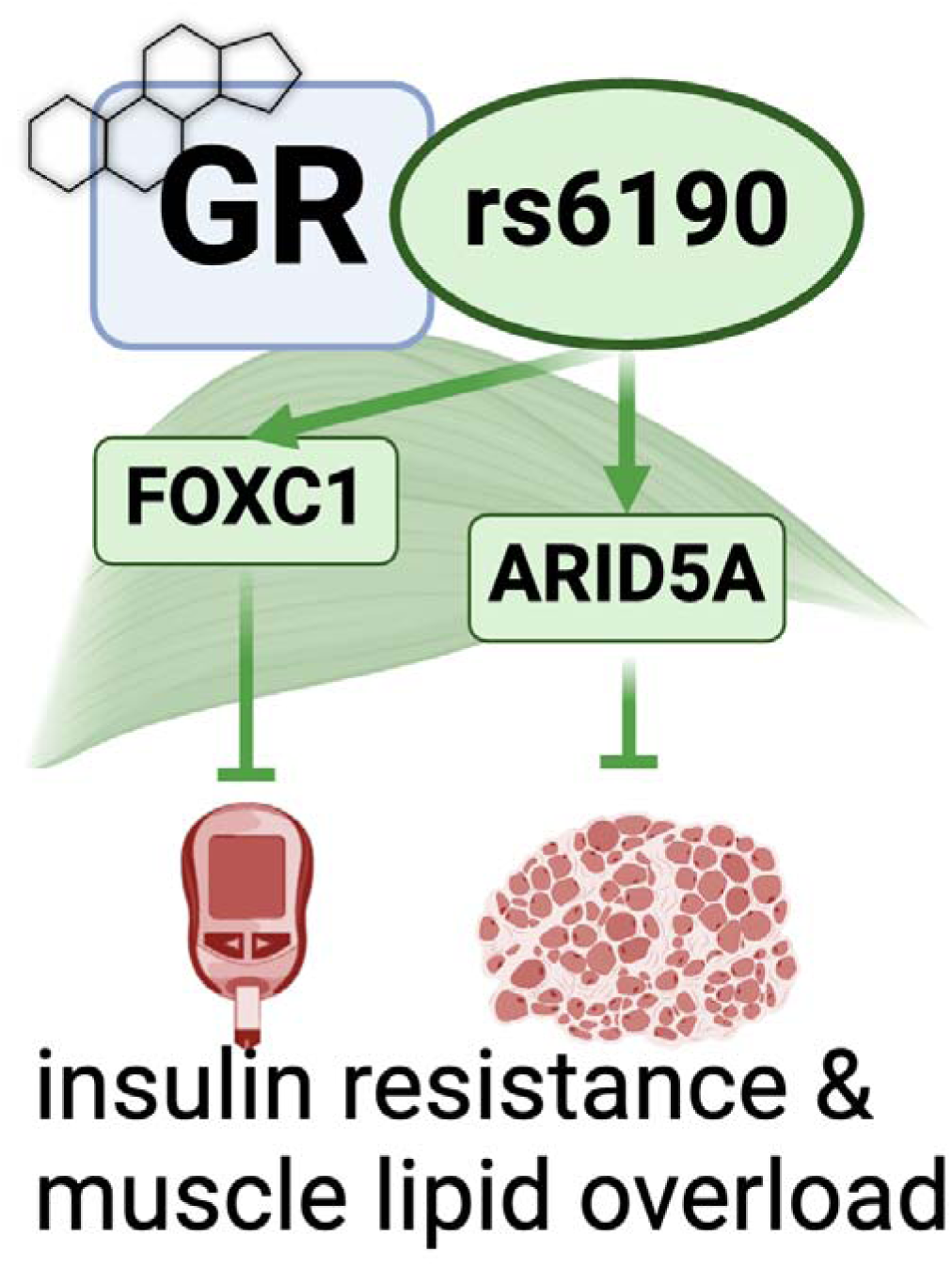

## Introduction

Insulin resistance is a well-known pathophysiological marker and risk factor for type 2 diabetes and cardiovascular diseases ^1^. Indeed, the insulin resistance observed in obese diabetic subjects directly impacts metabolic health through reduced systemic glucose clearance and progressive loss of lean muscle mass ^2,3^. Recent successes in glycemic control through antihyperglycemic drugs are positively impacting the management heart failure risk ^4,5^. However, muscle-centered mechanisms to rescue lean mass and strength in conditions of insulin resistance remain very limitedly elucidated. This becomes critical considering the sharp rise in prevalence of diabetes and obesity among adults, which will near about 50% of the global population by 2040 ^6,7^ and therefore create an urgent need to identify molecular targets that could remodel the insulin resistant muscle towards metabolic competence. Indeed, skeletal muscle is a major determinant (up to 80%) of insulin-mediated glucose disposal and utilization in both humans and rodents ^8,9^.

Glucocorticoids (GC) exert multiple pleotropic actions critical for metabolic, physiological, and stress-related conditions through activation of the glucocorticoid receptor (GR; *NR3C1* gene) ^10,11^. Glucocorticoids (GCs) play a crucial role in regulating metabolic homeostasis of glucose, lipid and protein in skeletal muscle development ^12–14^. The response of skeletal muscle to the GR action is modulated by single nucleotide polymorphisms (SNPs) that can impact metabolic homeostasis through a modified GR protein function ^15^. In humans, several SNPs within the 9 exons of the GR gene have been identified and studied for their association with glucocorticoid sensitivity and pathophysiological impact on human health ^16,17^. These genetic variations can affect the function, expression, or regulation of the glucocorticoid receptor, leading to differences in the individual response to glucocorticoid hormones ^18^. Intriguingly, some single nucleotide polymorphisms (SNPs) including Asn363Ser (rs6195) and BclI (rs41423247) are associated with enhanced exogenous and endogenous glucocorticoid sensitivity predisposing those carriers to metabolic dysfunction that includes increased BMI, low bone density, insulin resistance and altered cholesterol levels that promote cardiovascular risk ^19,20^. On the contrary, the rs6190 SNP (p. R23K; also known as ER22/23EK because in complete linkage with the silent E22E rs6189 SNP) correlated with enhanced muscle strength, lean body mass and metabolic health in men in limited human cohorts ^21–23^. However, genetic proof and mechanism of action for a direct effect of rs6190 on metabolic health are still missing.

To investigate the mechanism of this variant GR we generated transgenic mice genocopying the rs6190 SNP to test whether and how the SNP affects metabolism. Based on transcriptomic and epigenomic datasets from muscle, we further explored Forkhead box C1 (*Foxc1* gene) and AT-Rich Interaction Domain 5A (*Arid5A* gene) as novel muscle-autonomous transactivation targets determining the mutant GR action on metabolism and action. Further, we validated requirement and sufficiency for these two factors in insulin sensitivity and muscle lipid accumulation through AAV-based myocyte-specific overexpression. We further probed the large UK Biobank and All of Us datasets to query for the SNP effect on markers of glucose homeostasis and strength. Our study leverages a human SNP mechanism of action to identify novel myocyte-autonomous targets to salvage exercise tolerance and insulin resistance from metabolic stress.

## Results

### GR^R24K/R24K^ mice exhibit improved insulin resistance and exercise tolerance

In order to gain direct genetic and biological insight in understanding the role of the non-synonymous coding rs6190 SNP in the GR gene *NR3C1* (transcript ENST00000231509.3 (- strand); c.68G>A; p.R23K)^24^, we generated a transgenic mouse model where we CRISPR-knocked-in a single nucleotide mutation in the orthologous codon of the endogenous murine *Nr3c1* gene (NM_008173 transcript; c.71G>A, p.R24K; **Suppl. Fig. 1A-B**). We then compared homozygous mutant mice (GR^R24K/R24K^) to non-mutant littermates (GR^wt/wt^) to maximize the potential SNP effect and simplify the comparison through homogenous GR pools (100% mutant vs 100% non-mutant GR pools). Also, we focused our comparisons on young adult (4mo) male mice considering the seminal correlations of the SNP with metabolic health in young adult men ^25^. We compared mutant to WT mice after 12-week-long exposures to ad libitum feeding with regular chow vs 60% kcal high-fat diet to challenge body-wide and muscle metabolism with diet-induced obesity and insulin resistance.

GR^R24K/R24K^ mice showed a smaller but leaner body compared to GR^wt/wt^ littermates at 4 months of age, i.e. smaller weight with lower fat mass and higher lean mass, significantly reducing the diet-induced adverse effects on lean and fat mass **(Figure 1A-C; Suppl. Fig. 2C)**. We tested for exercise tolerance through work until exhaustion at a 15°-inclined treadmill, and muscle force at the in vivo hindlimb dorsiflexion assay ^26^. Compared to GR^wt/wt^, GR^R24K/R24K^ mice exhibited increased values of treadmill work and max force, rescuing those parameters to control-like levels after high-fat diet **(Figure 1D-E)**. We then tested the overall glucose homeostasis through hyperinsulinemic-euglycemic clamp in conscious unrestrained mice^27^, HOMA-IR ^28^, as well as glycemia measurements and muscle 2DG uptake assays ^29^. Compared to GR^wt/wt^, GR^R24K/R24K^ mice showed increased insulin sensitivity in both diets, as shown by increased glucose infusion rate (GIR) during the clamp and decreased HOMA-IR **(Figure 1F-G)**. In accordance with the trends in insulin sensitivity, insulin-driven 2DG uptake (performed at the end of the clamp) in muscle was increased **(Figure 1H)**. Considering the apparent increase in muscle insulin sensitivity, we further characterized muscles for myofiber typing, myofiber cross-sectional area and oxidative function. Compared to GR^wt/wt^, GR^R24K/R24K^ mice showed a gain of oxidative myofiber-associated myosins, as shown by immunostaining, WBs and qPCRs for Myh4 (type 2B), Myh2 (type 2A) and Myh7 (type 1) **(Suppl. Fig. 1D-F)**. Compared to GR^wt/wt^, GR^R24K/R24K^ muscle showed increased myofiber cross-sectional area, a parameter indicative of gained muscle mass **(Suppl. Fig. 1G)**. We also tested mitochondrial complex abundance and muscle glucose oxidation, a direct marker of muscle insulin sensitivity ^30^. Mitochondrial complexes showed non-significant upward trends in the GR^R24K/R24K^ muscle, with complex IV showing a significant gain **(Suppl. Fig. 1G)**. Glucose oxidation in muscle tissue was increased in GR^R24K/R24K^ muscle, as shown by basal respiration and calculated ATP production in glucose-fueled Seahorse assays using muscle tissue biopsies^31^ **(Suppl. Fig. 1I)**. The changes in muscle glucose metabolism were paralleled by improved glycemia in GR^R24K/R24K^ vs GR^wt/wt^ mice, particularly in the fed state **(Figure 1I)**. Collectively, these findings indicate that the rs6190 SNP is sufficient to protect insulin sensitivity and exercise tolerance against metabolic stress.

**Figure 1.**
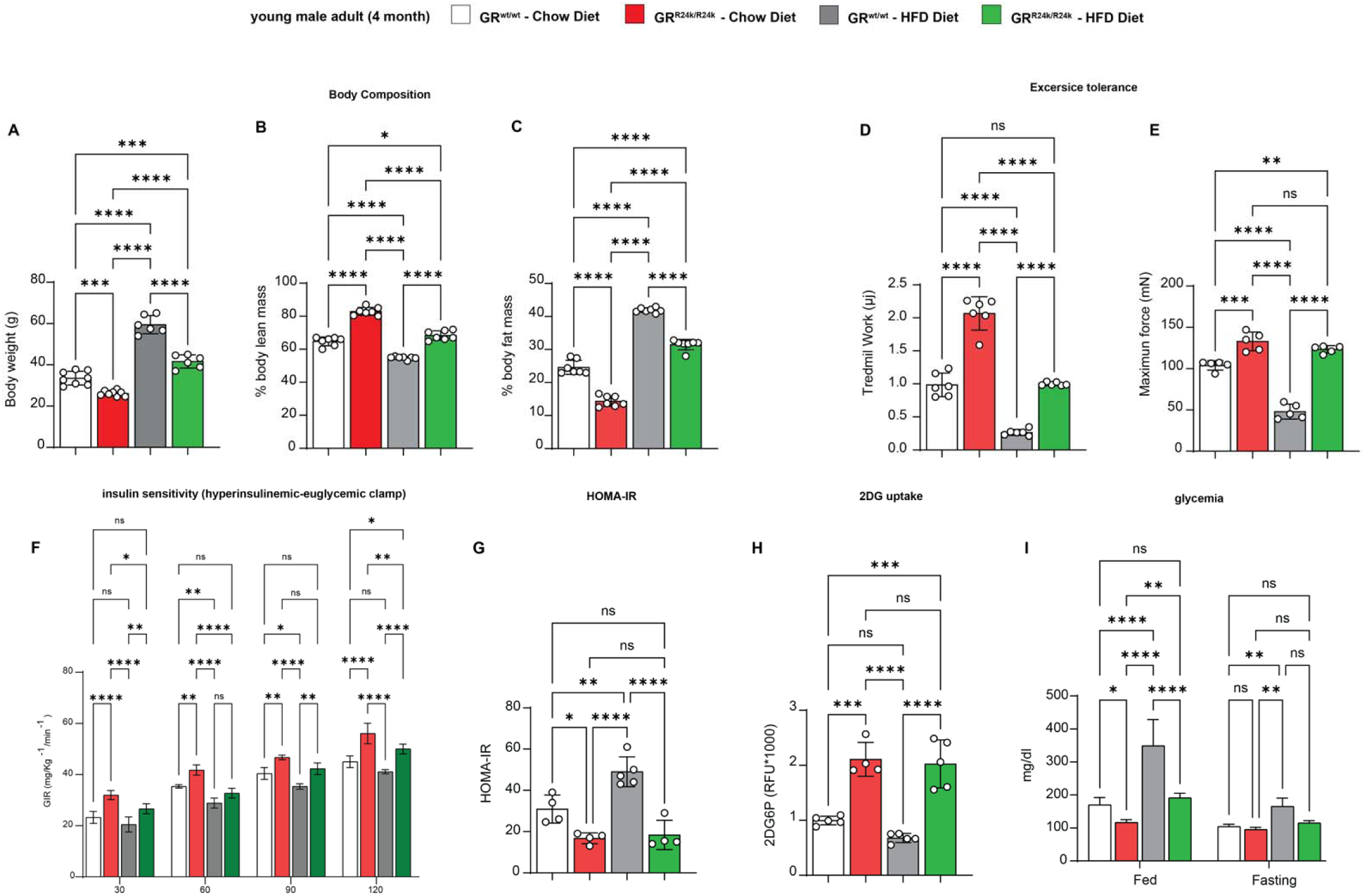
GR-R24K mice are protected from diet-induced obesity, insulin resistance and weakness. (A-. **C)** Compared to WT littermates, R24K-homozygous mice showed leaner body composition with normal and high-fat (60% kcal) diets. **(D-E)** The genetic variant increased aerobic capacity (treadmill work until exhaustion) and muscle force (hindlimb dorsiflexor assay) in normal diet, rescuing those parameters after high-fat diet feeding. **(F-G)** R24K-homozygous mice showed increased or rescued insulin sensitivity, as shown by increased glucose infusion rates (GIRs) during a hyperinsulinemic-euglycemic clamp and HOMA-IR values at endpoint of diet regimens. **(H)** Muscle insulin sensitivity and glucose uptake were increased by the R24K variant, as shown by muscle 2DG uptake at the end of the hyperinsulinemic-euglycemic clamp. **(I)** R24K-homozygous mice showed reduced or rescued glycemia in the fed state (ZT0, after feeding during the active phase). n=3-7♂/group; diet exposures for 12 weeks from 4mo to 7mo; 2w ANOVA + Sidak: ns, non significant; *, P<0.05; **, P<0.01; ***, P<0.001; ****, P<0.0001.

### In muscle, the mutant GR exhibits a specific transactivation program targeting Foxc1 and Arid5a

Considering the changes in muscle insulin sensitivity and metabolism, we focused on muscle to delve for molecular and epigenetic mechanisms of SNP action on the GR. The R24K mutation did not significantly change overall GR protein levels **(Figure 2A)**, despite a slight increase in mRNA expression **(Suppl. Fig. 2A)**. Because the amino acid substitution is in the N-terminal domain of the GR, which mediates protein-protein interactions ^32^, we sought to gain insight in potential changes in the GR interactions with other proteins in vivo. We performed an immunoprecipitation-mass spectrometry screening for GR interacting proteins in quadriceps muscles of GR^wt/wt^ vs GR^R24K/R24K^ mice. Strikingly, we found that the mutant GR displayed a strong downregulation in the binding of Hsp70 complex members **(Figure 2B)**, which we validated through CoIP **(Suppl. Fig. 2A)**. Because Hsp70 is a major cytoplasmic docker for the GR before its nuclear translocation ^33^, we tested the mutant GR translocation capacity in muscle comparing the GR protein signal in nuclear vs cytoplasmic fractions at 30min after a single dexamethasone injection. Compared to the WT GR, the mutant GR showed increased nuclear translocation capacity **(Figure 2C)**. Considering the skew in nuclear translocation, we tested the extent to which the SNP changed the epigenomic activity of the muscle GR through muscle GR ChIP-seq in quadriceps muscle. The GR binding element (GRE) motif was the top enriched motif in the datasets from both GR^wt/wt^ and GR^R24K/R24K^ muscles, as well as the typical expected GR peaks on the canonical GR reporter *Fkbp5* promoter were clearly defined **(Suppl. Fig. 2C)**, validating our datasets. Genome-wide occupancy on GRE motifs genomewide was increased by the mutant GR, as shown by density plot and heatmaps, although no genotype-related shifts in overall peak distribution (highly enriched in promoter-TSS regions) were observed **(Suppl. Fig. 2D-E)**. To find potential gene targets of the increased epigenomic activity of the mutant GR, we overlayed our ChIP-seq datasets with RNA-seq datasets that were obtained from subfractions of the same muscle samples. We ranked differentially expressed genes for mutant GR-dependent gains in GR peak signal in the promoter-TSS region and in overall RNA fold change, and we *found forkhead box C1* (*Foxc1* gene) and *AT-Rich Interaction Domain 5A (Arid5A* gene) as top hits **(Figure 2C)**. Both genes were upregulated at the protein level in GR^R24K/R24K^ vs GR^wt/wt^ muscle in normal and high-fat diet conditions **(Figure 2D)** and showed a clear gain of promoter-TSS GR peak in GR^R24K/R24K^ muscles **(Suppl. Fig. 2F)**.

**Figure 2.**
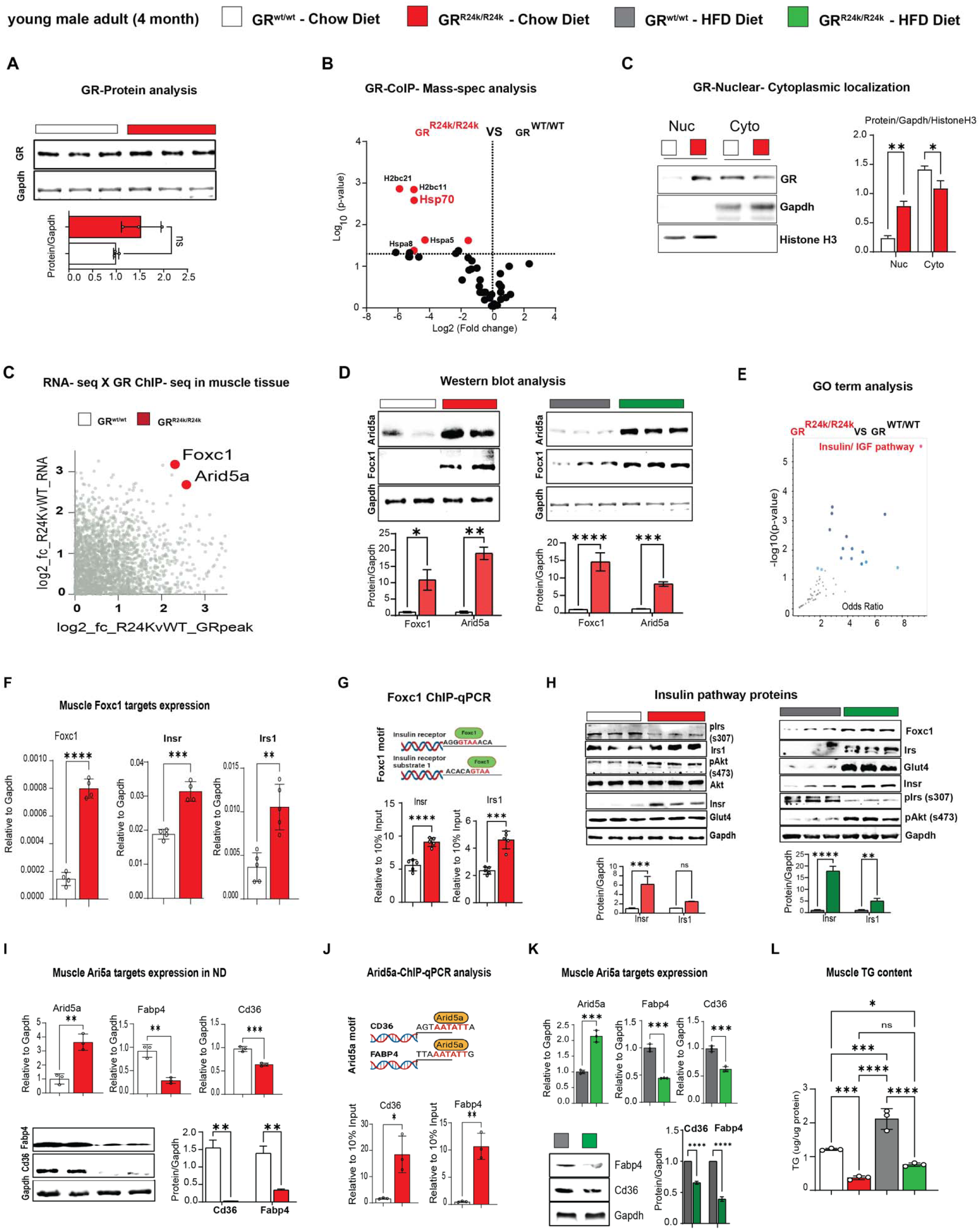
Foxc1 and Arid5a are novel transactivation targets of the mutant GR in muscle. (A-C) While the overall GR levels were not significantly changed by the R24K variant in muscle, the mutant GR decreased its interaction with proteins in the Hsp70 complex (IP-MS, red dots) and increased dexamethasone-induced nuclear GR translocation in muscle. **(C-D)** Overlay of RNA-seq and GR ChIP-seq in muscle unveiled two novel transactivation targets for the mutant GR, *Foxc1* and *Arid5a*, which were indeed increased in R24K vs WT muscle at the protein level. **(E)** The “insulin-IGF1 pathway” was enriched in an unbiased gene ontology analysis of the SNP-driven muscle transcriptomic changes, and included *Insr* and *Irs1*. **(F-H)** We identified Foxc1-binding elements in the proximal promoters of *Insr* and *Irs1*, which were indeed upregulated in the R24K vs WT muscle, along with Glut4 and p-AKT increases particularly after high-fat diet. **(I-L)** Arid5a transcriptional repression of Cd36 and Fabp4 increased in R24K vs WT muscle in both normal and high-fat diet conditions, lowering triacylglycerol accumulation in muscle. n=3-5♂/group; diet exposures for 12 weeks from 4mo to 7mo; Welch’s t-tests: ns, non significant; *, P<0.05; **, P<0.01; ***, P<0.001; ****, P<0.0001.

On one hand, Foxc1 is a Forkhead-box transcription factor implicated in Axenfeld-Rieger syndrome^34^ and kidney development ^35^, but never studied in muscle and metabolism. Cross-check through the predictive tool Harmonizome ^36^ unveiled that *Insr* (insulin receptor) and *Irs1* (insulin receptor substrate 1) genes were putative targets of *Foxc1* and were indeed the top upregulated genes in the enriched “insulin/IGF pathway” gene ontology term per GR^R24K/R24K^ vs GR^wt/wt^ RNA-seq comparison **(Figure 2E)**. *Foxc1*, *Insr* and *Irs1* upregulation in GR^R24K/R24K^ muscles was confirmed through qPCR, along with increased Foxc1 binding of canonical F-box sites in the proximal promoters of *Insr* (-103bp from TSS) and *Irs1* (-110bp from TSS) through ChIP-qPCR **(Figure 2F-G)**. Accordingly in the mutant muscle, upregulated Foxc1 protein levels correlated with increased Insr and Irs1 total levels, decreased inhibitory phosphorylation on Irs1-Ser307 (marker of IRS1 degradation in insulin resistant muscle ^37^) and increased levels of Glut4 and phosphorylation of AKT-Ser473, markers of insulin responsiveness ^38^, trends that were particularly reinforced in the obese muscle **(Figure 2H)**. In vitro, *Foxc1* overexpression through C2C12 myoblast transfection increased the total protein levels of Insr and Irs1 **(Suppl. Fig. 2G).**

On the other hand, *Arid5A* has been reported in adipose tissue as pro-metabolic factor by repression of lipid transport genes *Cd36* and *Fabp4* expression ^39^, but its function in muscle remains virtually unknown. Compared to GR^wt/wt^, the *Arid5A* upregulation in GR^R24K/R24K^ muscle correlated with downregulation of *Cd36* and *Fabp4* levels, which in turn showed increased occupancy of Arid5A on their gene promoter sites (-745bp for *Cd36* TSS,-750bp for *Fabp4* TSS; **Figure 2I-J)**. The trends in Arid5A, Cd36 and Fabp4 expression and protein levels were replicated in the obese muscle **(Figure 2K)**. In line with the reported role of *Arid5a* in limiting lipid uptake and storage in adipose tissue ^39^, compared to GR^wt/wt^ the GR^R24K/R24K^ muscle showed lower levels of muscle triacylglycerol accumulation **(Figure 2L)**. In vitro in C2C12 myoblasts, Arid5A overexpression reduced Cd36 and Fabp4 levels **(Suppl. Fig. 2H)**. In silico prediction through STRING ^40^ suggested possible interaction of Arid5A with the repressor SAP30, a component of the repressive histone deacetylation complex that includes HDAC1 and SIN3A ^41^. We tested these protein interactors through CoIP and found that the mutant muscle showed increased recruitment of SAP30, HDAC1 and SIN3A proteins by Arid5A **(Suppl. Fig. 2I-J)**.

Taken together, these data show that in muscle the SNP increases a pro-metabolic epigenetic GR program, characterized by transactivation of *Foxc1* and *Arid5a*, whose causative roles in muscle remain unknown.

### Foxc1 and Arid5a are required and sufficient for muscle insulin sensitivity and lipid burden regulation

We sought to test rigorous genetic proofs of causal roles for *Foxc1* and *Arid5a* in the putative gene programs suggested by the mutant muscles. To test *Foxc1* sufficiency in muscle in vivo, we generated AAVs to overexpress either GFP (control) or *Foxc1* downstream of a *CMV* promoter. A strong adult myocyte tropism was promoted by using the MyoAAV serotype ^42^. Widespread transduction of muscles in vivo was confirmed through GFP **(Figure 3A)**. At 4 weeks after a single r.o. injection of 10^12^vg/mouse AAV vectors in WT mice, we found that *Foxc1* overexpression increased Insr and Irs1 levels, as well as muscle 2DG uptake **(Figure 3B-C)**. Analogously, MyoAAV-Arid5a was sufficient to repress Cd36 and Fabp4 in muscle, along with triacylglycerol levels **(Figure 3D-F)**. We then tested the extent to which the concerted increase in *Foxc1* and *Arid5a* in muscle was sufficient to mimic the SNP metabolic protective effect with high-fat diet. We injected WT mice with the combination of MyoAAV-Foxc1 and MyoAAV-Arid5a, using the control vector (GFP) as control, and then exposed them to the same 12-week-long high-fat diet. Overexpression of both factors in muscle recapitulated the molecular effects of each factor (Insr and Irs1 gain for Foxc1; Cd36 and Fabp4 loss for Arid5A) and resulted in improved glucose homeostasis and muscle lipid accumulation, as shown by reduced fasting glycemia, increased muscle 2DG uptake and reduced muscle triacylglycerols **(Figure 3G)**. As novel transactivation targets of the muscle GR, we tested whether the prednisone regimens we reported with opposite metabolic effects in muscle^43^ had also divergent effects on Foxc1 and Arid5a. We compared the Foxc1 and Arid5a cascades in muscle after 12-week-long intermittent (once-weekly, insulin-sensitizing effect) vs chronic (once-daily, insulin-desensitizing effect) in parallel in normal vs high-fat diet conditions. In both diets, intermittent prednisone increased Foxc1 and Arid5a programs in muscle, while daily prednisone decreased them **(Figure 3H)**. Finally, we sought proofs of requirement through MyoAAV-based, shRNA-driven knockdown of *Foxc1* or *Arid5A*, a strategy we also recently reported in adult obese diabetic heart in vivo^44^. *Foxc1* knockdown in muscle in vivo decreased Insr and Irs1 protein levels in muscle and, accordingly, 2DG uptake **(Figure 3I)**. *Arid5a* knockdown de-repressed Cd36 and Fabp4 protein levels, promoting triacylglycerol accumulation **(Figure 3J)**. Taken together, these data indicate that the myocyte-autonomous Foxc1-Arid5a program is required and sufficient to protect muscle insulin sensitivity and lipid burden from metabolic stressors, and appears a “linchpin” axis in determining the pro-vs anti-metabolic outcome of glucocorticoid stimulation.

**Figure 3.**
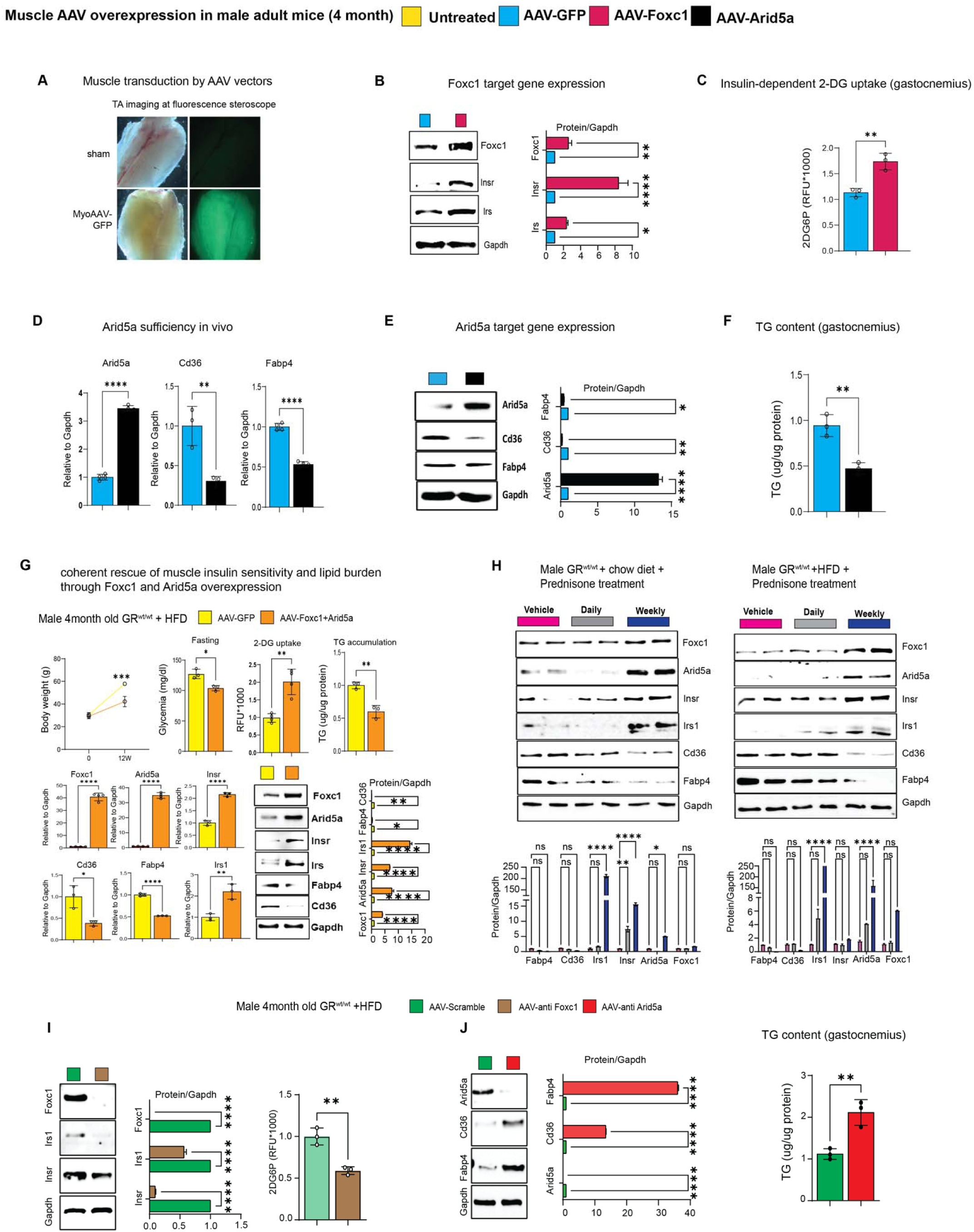
Foxc1 and Arid5a directly regulate muscle insulin sensitivity and lipotoxicity. **(A)** Muscle-wide transduction by the MyoAAV-serotyped vectors. **(B-C)** In WT mice on normal diet, *Foxc1* overexpression is sufficient to upregulate Insr and Irs1, and increase insulin-driven 2DG uptake in muscle. **(D-F)** In WT mice on normal diet, *Arid5a* overexpression is sufficient to lower Cd36, Fabp4 and triacylglycerol (TG) levels in muscle. **(G)** Co-transduction of MyoAAV-Foxc1 and-Arid5a reduces the effects of diet-induced obesity on glycemia, muscle glucose uptake and muscle TG content, while recapitulating each of the target gene effects for either Foxc1 or Arid5a. **(H)** Foxc1 and Arid5a gene programs in muscle are upregulated by the insulin-sensitizing once-weekly prednisone treatment, while downregulated by the insulin-desensitizing once-daily prednisone treatment. **(I-J)** MyoAAV-driven knockdown in muscle in high-fat diet conditions showed Foxc1 requirement for *Insr, Irs1* expression and muscle 2DG uptake, and Arid5a requirement for *Cd36*, *Fabp4* repression and muscle TG lowering. n=3-5♂/group; diet exposures for 12 weeks from 4mo to 7mo; Welch’s t-tests and 1w ANOVA + Sidak (H): ns, non significant; *, P<0.05; **, P<0.01; ***, P<0.001; ****, P<0.0001.

### Data from the UK Biobank support a pro-metabolic effect of the mutant GR in humans

To gain further insight in the relevance of our SNP-related metabolic mechanism of action for humans, we probed the large dataset of the UK Biobank that comprises data from 485,895 adults of ∼40-70 years of age. In this cohort, the GR rs6190 variant (*NR3C1* gene, transcript ENST00000231509.3 (-strand); c.68G>A; p.R23K) exhibited a minor allele frequency of 2.68%, with 25,944 heterozygous individuals and 413 homozygous individuals for the rs6190 SNP. We stratified parameters of relevance aligned with our prior measures of metabolic and muscle function, i.e. body mass index (BMI), glycemia, lean mass and hand grip strength normalized to arm lean mass, according to rs6190 SNP zygosity. We are defining here homozygous carriers of the reference allele (control population) as GR^ref/ref^, heterozygous SNP carriers as GR^ref/ALT^, and homozygous SNP carriers as GR^ALT/ALT^. We performed linear regressions with a mixed model correcting for age, diabetic status, lipidemia levels and top 10 principal components. Consistent with prior associations in heterozygous carriers in small cohorts ^25^, we found significant correlations of rs6190 SNP in sex-aggregated UK Biobank population with decreases in BMI, glycemia and increases in lean mass and hand grip strength according to zygosity **(Figure 4A)**. To probe these rs6190 correlations in a more genetically diverse human dataset, we queried the All Of Us dataset, where we found the SNP at a variable minor allele frequency ranging from low-frequency to rare across ancestries: African/African-American, 0.49%; American Admixed/Latino, 0.84%; East Asian, 0.061%; European, 2.67%; Middle Eastern, 1.43%; South Asian, 1.49%. In the All Of Us subset of 245,385 individuals with rs6190 genotype annotation encompassing all ancestries and ages, we repeated the linear regressions corrected for age, diabetes, lipidemia and top ten principal components. The regressions in the All of Us dataset confirmed a significant correlation between rs6190 zygosity and declining trends in BMI, glycemia, insulinemia and A1C **(Figure 4B)**, all parameters typically related to glucose homeostasis and metabolic health in humans. Taken together with our genetic studies in mice, these data further support the potential relevance of the pro-metabolic mechanisms enabled by the rs6190-mutant GR for human metabolic health.

**Figure 4.**
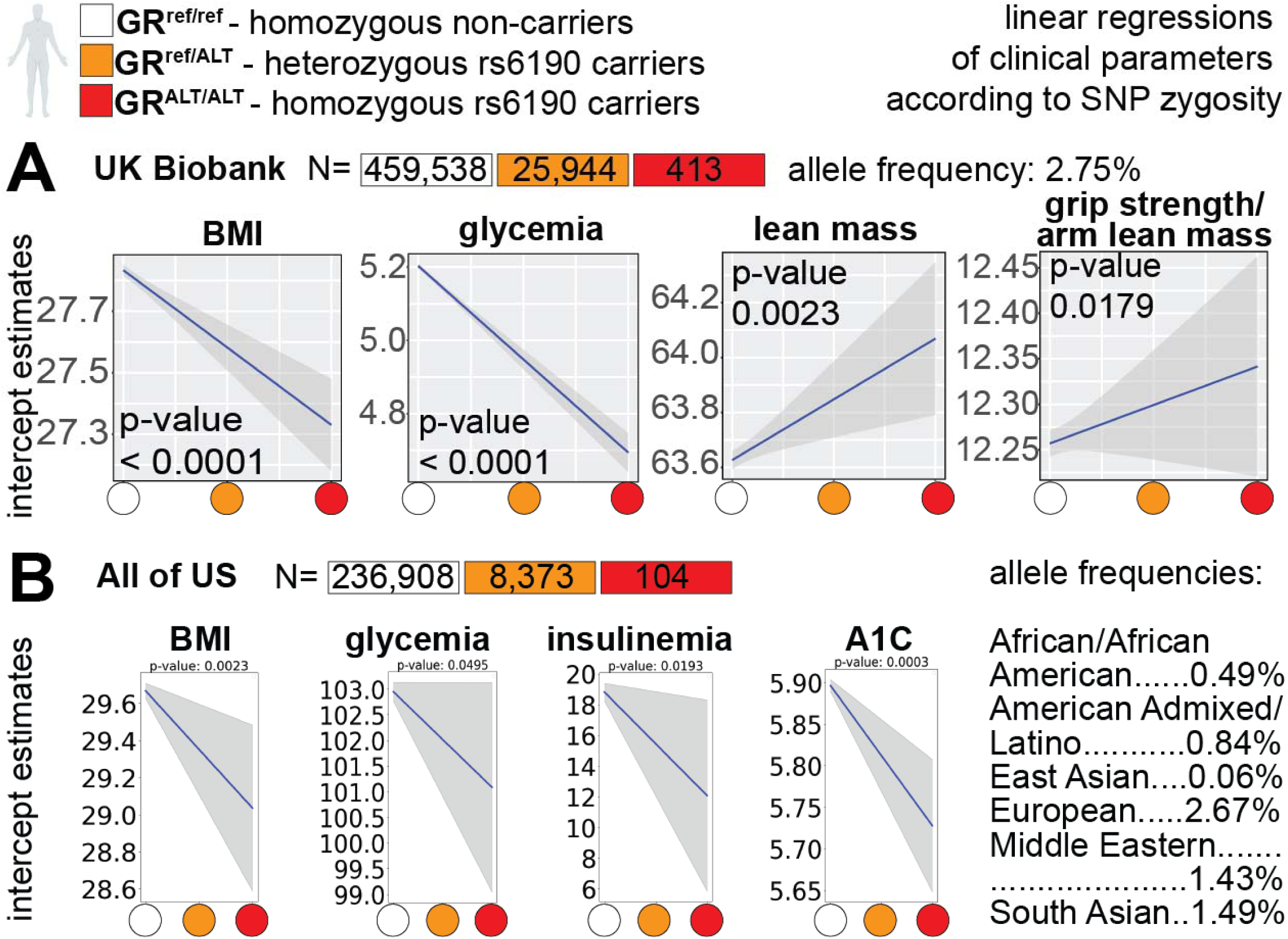
Genotype for the mutant GR (rs6190 SNP) correlates with metabolic health in humans –. **(A)** rs6190 zygosity correlated with decreases in BMI and glycemia and increases in lean mass and grip strength in the UK Biobank. **(B)** SNP zygosity correlated with decreases in BMI, glycemia, insulinemia and A1C in the All of US (ancestry-segregated allele frequencies are reported). Data for both cohorts is shown aggregated for sex, ages and ancestries.

## Discussion

Muscle insulin sensitivity is critical for metabolic health ^8^, as skeletal muscle accounts for ∼80% of glucose uptake postprandial ^45^ or after an oral bolus ^46^. The insulin-resistant muscle progressively loses mass and function, exacerbating the vicious circle of metabolic stress and exercise intolerance ^47^. Indeed, in type-2 diabetes, muscle insulin resistance generally precedes beta cell failure and overt hyperglycemia ^48^. However, the quest for actionable muscle-autonomous mechanisms to rescue insulin sensitivity is still open. Here we leverage the mechanism of action of the rs6190-mutant GR in muscle to unveil the potentially critical role of *Foxc1* and *Arid5a* as unprecedented muscle-autonomous factors sufficient to promote overall insulin sensitivity and reduce lipotoxicity. Our in vivo sufficiency/requirement proofs through myotropic AAVs indicate their causal role in the muscle response to the challenges posed by high-fat diet on glucose homeostasis and insulin sensitivity. It must also be noted that our study is the first to report myocyte-autonomous roles and molecular targets for both Foxc1 and Arid5a in muscle, paving the way to future studies delving in those cascades.

Glucocorticoid steroids and the glucocorticoid receptor (GR) constitute a primal circadian axis regulating glucose homeostasis and insulin sensitivity, as evidenced by the prefix “gluco” and the long-known effects on liver gluconeogenesis^49^ and adipose tissue lipolysis^50^. Glucocorticoids are widely prescribed to manage inflammation and are used by over 2.5mln people for over 4 years in the US alone ^51^. Typically, glucocorticoids are prescribed to be taken once-daily at the start of our active-phase (early morning) ^52^, but such glucocorticoid regimens are very well known to disrupt insulin sensitivity, particularly in muscle ^12^. Recently, we have discovered that intermittence in chronic frequency-of-intake ^43^ and early rest-phase as circadian time-of-intake ^53^ uncover pro-ergogenic glucocorticoid-GR mechanisms in muscle. In that regard, GR mechanisms of insulin sensitization are emerging in non-muscle cells, from the adipocyte GR stimulating adiponectin ^43^ to the macrophage GR protecting against insulin resistance ^54^. However, myocyte-autonomous mechanisms of insulin sensitization by the glucocorticoid receptor remain quite unanticipated in the field. Here we report that a non-synonymous human variant in the glucocorticoid receptor skews its activity towards a pro-metabolic program in muscle. The divergent trends in Foxc1 and Arid5a programs in intermittent (pro-metabolic) vs daily (anti-metabolic) prednisone treatments further corroborate the discovery of a “linchpin” axis discriminating between pro-and anti-metabolic outcomes of the GR activation in muscle. Our muscle-centered study is the first in vivo investigation of the potential physiologic mechanisms enabled by the rs6190-mutant GR, and future studies in other tissues of metabolic interest will help articulate an holistic paradigm for the metabolic impact of this non-rare mutant GR in the human population.

We are focusing on Foxc1 and Arid5A as putative muscle GR effectors of insulin sensitivity and lipotoxicity protection thanks to a rather uncommon angle, i.e. the human GR variant rs6190. Traditionally, rs6190 is referred to as ER22/23EK or rs6189/rs6190 due to the complete linkage with the silent rs6189 SNP on the previous codon (E->E). The rs6190 SNP case is fascinating because a theoretically inconsequential coding variant (conservative replacement R->K in position 23) associated with lower levels of fasting insulin and HOMA-IR ^22^, increased lean mass and muscle strength as young adults ^25^, and prolonged survival as older adults ^55^. The proposed mechanism of “glucocorticoid resistance”, based on limited in vitro observations ^56^, was largely unreproducible in many other association studies ^24,57–62^. Thus, the extent to which the coding rs6190 GR variant is sufficient to directly regulate insulin sensitivity, as well as the underlying mechanism, remain unknown. We re-assessed the rs6190 associations in the large UK Biobank and All of Us datasets, and tested sufficiency and mechanism for the SNP in CRISPR-engineered mice, confirming that the SNP is sufficient to increase muscle insulin sensitivity. However, in contrast to previously proposed “glucocorticoid resistance”, we found that in muscle tissue the SNP *increased* the dexamethasone-induced nuclear translocation (GR activation), as well as its epigenomic activity. Also, we found Foxc1 and Arid5A as consequential targets of mutant GR-driven transactivation in muscle. Therefore, taken together, our data challenge the paradigm of this mutation on GR activity at least in muscle, and open a compelling avenue of investigation in other systems of relevance, like liver, adipose tissue and immune system.

## Acknowledgements

Mass-spec analyses were performed thanks to the Proteomics Mass-Spec Core Facility at University of Cincinnati, with critical assistance by Dr. Greis and Dr. Haffey. Next-generation sequencing was performed thanks to the Cincinnati Children’s DNA Sequencing and Genotyping Facility (RRID: SCR_022630), with critical assistance by David Fletcher, Keely Icardi, Julia Flynn, and Taliesin Lenhart.

## Materials and Methods

### Mice handling and Transgenic mice generation

Mice handling and maintenance in polypropylene cages with chow diet and water ad libitum were done as per the American Veterinary Medical Association (AVMA) and under protocols fully approved by the Institutional Animal Care and Use Committee (IACUC) at Cincinnati Children’s Hospital Medical Center (#2022-0020, #2023-0002). Mice, which is a well-established model system for metabolic research were maintained in a controlled room temperature of @22°C with 14/10 hr light/dark cycle in a purpose build pathogen free animal facility consistent with the ethical approval. Periodic change of cages, with fresh water and beds, was done to ensure a healthy and stress-free environment for the animals. Rodent diet with 60 kcal% fat (Research Diets, D12492) was used to generate High Fat Diet induced obese (HFD) animal groups.

WT mice were obtained and interbred from the Jakson Laboratories (Bar Harbor, ME; JAX strain) as WT C57BL/6 mice #000664. Transgenic mice genocopying the polymorphism R24K was established through the CRISPR/Cas9 genome editing in the endogenous *Nr3c1* locus on the C57BL/6J background. This genetic modification was performed by the Transgenic Animal and Genome Editing Core Facility at CCHMC. To ensure genetic background homogeneity and control for potential confounding variables, the colonies were maintained through heterozygous mattings. This approach allowed us to compare two distinct groups of male mice as littermates: GR^wt/wt^ (control WT) and GR^R24K/R24K^ (homozygous SNP carriers) in homogenous genetic background conditions.

### DNA isolation and Genotyping

DNA isolation from tail/ muscle tissue for genotyping experiments were done using the kit from G biosciences (#786-136). Briefly, Samples (ear, toe, tail and muscle tissue) were collected in a 1.5ml micro centrifuge tube containing 500ul of genomic lysis buffer and 10ul of proteinase K solution incubated on thermomixer at 60C for 3-4 hrs or overnight. The samples were cooled to room temperature and 200ul of chloroform were added and mixed by inverting several times centrifuged at 14000g for 10 minutes. The upper phase was separated to a new clean 1.5 ml micro centrifuge tube and 150ul of precipitation solution were added and centrifuged for 5 min at 14000g. Transfer the supernatant to a new 1.5 ml micro centrifuge tube and add 500ul isopropanol invert it several times and centrifuge at 14000g for 5 min to precipitate the genomic DNA. Add 700ul of 70% ethanol to wash the DNA pellet and centrifuge for 1min at 14000g. decant the ethanol and air dry the pellet for 5 min or until no ethanol is observed. Add 50ul of MilliQ water to the DNA pellet and incubate in the thermomixer at 55C for 15 min to rehydrate or at 4C in fridge O/N.

Genotyping the R24K mice carrying the GR^wt/wt^ / GR^R24K/R24k^ polymorphism were genotyped by PCR-RFLP method. Briefly, 18 µl of PCR master mix which includes MM (Promega #xxx), 1ul of Forward/Reverse primers (10mM), nuclease free water and 2 µl of the isolated DNA were subjected to Polymerase chain reaction with the 40 cycles (95C-10 min and 40 cycle of 95C-30sec, 55C-30sec, 72C-30sec and final 72C-5min). After the PCR 20ul of the PCR product were restriction digested with BamH1 (NEB #xxx) for 1 hr at 37C. The digested PCR product was resolved on 2% Agarose gel and visualized in a UV transilluminator. The mice genotypes were denoted based on their band size (GR^wt/wt^-xbp and GR^R24K/R24K^-xbp). Primers used for the genotyping; for-TGTACATTTAGCGAGTGGCAGGAT; Rev-TGCTGAGCCTTTTGAAAAATCAAG; GR wild type has a band size of 474bp while the GR R24K-14bp.

### Tissue and blood collection, assessment of glucose and insulin tolerance test and 2-DG uptake, Triglyceride estimation

All young adult mice (4-month-old) used for the experiments were euthanized through carbon dioxide inhalation followed by cervical dislocation and tissues of metabolic relevance such as skeletal muscle (soleus and gastrocnemius) was dissected out using a sterile surgical kit, rapidly snap frozen in liquid nitrogen and stored at-80°C for further analysis.

Blood collections were carried out by tail snip or euthanasia method. For the tail snip method, pups or the rats were restrained in the cage with the lid closed having the tail outside, and one or 2 mm of the tail tip was quickly cut using sterilized surgical scissors. By gently squeezing the tail from the base, the blood was collected and assessed for hyperglycemia using a glucometer (OneTouch® Ultra® 2 meter). For serum collection, the animals were euthanized, and around 1 to 2 ml of blood withdrawn using a sterile syringe from the abdominal aorta (unlock 3 ml syringe,). The blood was allowed to stand at room temperature, centrifuged at 5000 rpm for 5 min and the serum was transferred to a new tube and stored at-80°C for further analysis.

To perform glucose tolerance test on mice fed on standard chow and HFD diet, were fasted for overnight (12- 16hrs). The fasting glucose levels were assessed by tail snip method as described above using a glucometer as described. After the fasting glucose assessment, an intraperitoneal injection of D-glucose solution (Sigma, G8270) was injected at a rate of 2 g/kg body weight concentration to all the groups. The blood glucose levels were assessed by tail snip method at every 30 min till 2 hours post glucose injection and recorded. mice were then sacrificed, and the tissues and serum samples were collected as above. The same procedure was followed for insulin tolerance test with the fasted mice after glucose measurement at baseline injected with 0.5U/Kg insulin in 100ul PBS. Glycemia was recorded every 30 min post injection. The HOMA-IR calculation for analyzing the insulin resistance was also undertaken^63^.

The 2-DG glucose uptake for tissues was analyzed by Promega Glucose uptake-Glo assay method (#J1341). Briefly, 1mM solution of 2DG was injected into the mice 30 min before euthanasia. Tissue such as skeletal muscle was collected and crushed into fine powder and 20-50mg was used for the assay. Thaw all the reagents at room temperature and mix 25ul of neutralization buffer to 100ul of the reaction mixture per reaction to the powdered tissue. Mix and let it sit for 0.5- 5 hours and centrifuge of 5 min at 10000g. Separate 125µl of the supernatant into 96 well plate and read luminescence on a plate reader. In the same way the triglyceride accumulation and insulin were quantitated by kit method as per manufacturer’s protocol (CYMAN chemical # 10010303; 589501).

### RNA isolation, RNA- Sequencing, cDNA synthesis and Qualitative PCR analysis

RNA isolation was carried out by the standard Trizol method ^64^. The tissue sample was cut into pieces using a sterile blade and transferred to a 2 ml Eppendorf tube to which 1ml of Trizol reagent (Invitrogen # 15596018) was added. The tissue was homogenized using a Tissue lyzer (Benchmark, D1000) and 0.2 ml of chloroform was added and vortexed. The sample was centrifuged at 14000 rpm for 15 min which formed 3 different layers. The upper aqueous layer which contains the RNA, is separated into a new 1.5 ml Eppendorf tube and 500 µl of isopropyl alcohol was added, mixed by inverting and centrifuged at 14000 rpm for 10 min. After centrifugation, the pellet was washed with 70% ethanol and centrifuged at 14000 rpm for 10 min. Purification of RNA was carried out by adding 200 µl of DNase 1 buffer containing 5 µl of DNase 1. The pellet was reconstituted and incubated at 370C for 30 min. After incubation, 200 µl of lysis buffer and 200 µl of MPC solution was added, vortexed and incubated in ice for 5 min. Following incubation, the mixture was vortexed and centrifuged at 14000 rpm for 10 min. This step precipitates the proteins and salts leaving the upper aqueous layer containing the RNA which was separated carefully into a new tube and 500 µl of isopropanol were added and the tube was inverted several times and centrifuged at 14000 rpm for 10min to pellet the RNA. The pellet was washed using 70% ethanol by adding 500ul to the tube and centrifuged at 14000 rpm for 10 min. The tube was air-dried at room temperature and the pelleted RNA was re-suspended with 30 µl of milliQ water.

RNA-seq was performed at the DNA Core at the CCHMC facility with 10 ng – 150 ng of total RNA used after quantification by Qubit RNA HS assay kit (Cat #Q32852; Invitrogen, Waltham, MA). Based RNA integrity value above 7 determined by the spectrofluorometric measurement RNA samples was poly-A selected and reverse transcribed using Illumina’s TruSeq stranded mRNA library preparation kit (Cat# 20020595; Illumina, San Diego, CA). Library preparation was done for each sample fitted with one of 96 adapters with different 8 base molecular barcode for high level multiplexing and following 15 cycles of PCR amplification, completed libraries were sequenced on an Illumina NovaSeqTM 6000, generating 20 million or more high quality 100 base long paired end reads per sample. A quality control check on the fastq files was performed using Fast QC. Upon passing basic quality metrics, the reads were trimmed to remove adapters and low-quality reads using default parameters in Trimmomatic [Version 0.33]. In the next step, transcript/gene abundance was determined using kallisto [Version 0.43.1]. The trimmed reads were then mapped to mm10 reference genome using default parameters with strandness (R for single-end and RF for paired-end) option in Hisat2 [Version 2.0.5]. In the next step, transcript/gene abundance was determined using kallisto [Version 0.43.1]. We first created a transcriptome index in kallisto using Ensembl cDNA sequences for the reference genome. This index was then used to quantify transcript abundance in raw counts and counts per million (CPM). Differential expression (DE genes, FDR<0.05) was quantitated through DESeq2. PCA was conducted using ClustVis. Gene ontology pathway enrichment was conducted using the Gene Ontology analysis tool.

The conversion of RNA to cDNA was carried out with Superscript IV Vilo kit using 1µg of total RNA in a reaction volume of 20 µl as per manufacturer’s instructions (Invitrogen #11766050). The reaction mixture in 4 µl consisting of Mgcl2, dNTP mix, Random primer and Reverse Transcriptase was set up with the remaining 16 µl with 1 ug of RNA with nuclease free water. The reverse transcription was carried out in the thermal cycler with the following steps, i.e., 25°C for 10 min, 55°C for 10 min, 85°C for 5 min and hold at 4°C. The 20 µl reaction mixture was then reconstituted with Milli-Q water to 50 µl and used for further analysis. Quantitative RT-PCR reactions were carried out in a volume of 20 µl of 1X SYBR Green fast qPCR Mix (#RK21200, ABclonal, Woburn, MA), and 100mM primers using CFX96 qPCR machine (Bio-Rad, Hercules, CA; thermal profile: 95C, 15sec; 60C, 30sec; 40X; melting curve). Comparative C(T) method which is also referred to as the 2- ΔΔCT method ^65^ was used to determine the relative gene expression between the gene of interest relative to the internal housekeeping control gene. The internal control gene used in the assays was GAPDH. Primers used for the analysis are listed in Table1.

### Chromatin immunoprecipitation and sequencing

Chromatin Immunoprecipitation (ChIP) was carried out using the skeletal muscle for the transcriptomic analysis. The samples were chopped into small pieces and transferred to a tube containing 1ml of PBS with 27 µl of 37% formaldehyde. The cross-linking process was carried out for 10 minutes on a rotator. After the incubation 50 µl of 2.5 M glycine was added to each sample to a final concentration of 0.125 M and incubated for another 5 minutes to stop the cross-linking process. The samples were then centrifuged at 5000 rpm for 5 min to collect and the supernatant was discarded without disturbing the pellet. The pellet was then washed by suspending in ice-cold PBS and centrifuged at 5000 rpm for 5 minutes. This washing procedure was carried out three times. The pellet was suspended in 1 ml of FA lysis buffer (50 mM HEPES, 140 mM NaCl, 1 mM ETDA, 1% Triton x-100, 0.1% sodium deoxy cholate) containing protease inhibitor cocktail and 20% SDS and subjected to sonication. Sonication of the chromatin fragmentation was performed using Bioruptor (Diagenode, Liège, Belgium) with 45 on/off cycle for 10 minutes. After the sonication, the samples were centrifuged for 10 min at 14000 rpm to collect the supernatant/ lysate in a new tube. About 180 µl of the lysate was used for the immunoprecipitation (IP) with the specific antibody listed in Table 2. Twenty percent of the IP was taken as input and stored separately at-80°C for further use. The immunoprecipitation reaction of 500 µl consisting of the FA lysis buffer with the protease inhibitor cocktail and the lysate was used for each sample. The respective antibodies used are given in table 6 and an antibody concentration of 5 ug per sample was used. Pierce A/G magnetic beads (Invitrogen # 80105G) of 30 µl were washed using FA lysis buffer 2 times and mixed with the IP samples. The Immunoprecipitation reaction was carried out overnight on a rotator at 4 °C. After the incubation, the beads were separated using the magnetic stand and the other lysate was discarded. The lysate was washed with FA lysis buffer, high salt solution buffer, LiCl buffer and finally with TE buffer. The final elution was carried out by suspending the beads in 100 µl of elution buffer and incubation on a shaking dry bath for 10 minutes at 70°C. The bead was separated on the magnetic stand and the renaming elution buffer containing the protein DNA complex was collected in a separate tube. The input stored at-80 was used along with the IP samples. 4 µl of 5 M NaCl added to all samples and the reverse crosslinking were performed at 65°C overnight on shaking dry bath. Following that, the DNA isolation was carried out as described in the DNA isolation protocol. Percentage of Input, control and experimental samples were measured by qPCR analysis as described earlier. Primers were selected among validated primer sets from the MGH Primer Bank; IDs: INS- 117606344c1; MYH7- 18859641a1; MYH4- 9581821a1; MYH2-21489941a1; GR-6680103a1; Foxc1-410056a1; Arid5a-31542476a1; INSR-67543660a1;IRS-29568118a1; GAPDH-6679937a1; PPARG-6755138a1; CEBPA-6680916a1; FATP1-6755546a1; FABP4-14149635a1; CD36-31982474a1; ChIP-qPCR primers were manually designed using primer 3 software: INSR F- ACCGCCACTACTTCTGCTAC; INSR R- CTTGGATCTAGGCCCGTGG; IRS F- AAGGGGAGCAGGAGAAAAGG;IRS R-ACAAAAGGAGAACAGGGATCC;FABP4 F- CTGTAGCCCGCATCCAGAG;FABP4 R- TTGGCTTTGTTTGGTTTGGG; CD36 F- TAACCACCACAGCCATGAGT; CD36 R- CCACTTGGGGAAGCTGTTAG

For the ChIP sequence analysis, DNA purification with mini elute kit (Cat# 28004, QIAGEN, Hilden, Germany) following quantification using Qubit ds DNA quantification assay kit (Invitrogen #Q32851) was done and DNA concentration of 1ng was taken for analysis. Library preparation and sequencing were conducted at the CCHMC Genomics Core, using TruSeq ChIP-seq library prep (with size exclusion) on ∼10 ng of chromatin per ChIP sample or pooled inputs and HiSeq 50-bp was conducted using HOMER software (v4.10) after aligning fastq files to the mm10 mouse genome using bowtie2. PCA was conducted using ClustVis. Heatmaps of peak density were imaged with TreeView3. Peak tracks were imaged through WashU epigenome browser. Gene ontology pathway enrichment was conducted using the gene ontology analysis tool.

### Total protein isolation, Western blotting and Co-immunoprecipitation

Total protein isolation was carried out from skeletal muscle (soleus and gastrocnemius) tissues. About 100 mg of tissues were weighed and chopped into small pieces using a sterile blade and transferred to a 2 ml sterile Eppendorf tube. To each sample, 1 ml of RIPA lysis buffer was added. The RIPA buffer preparation includes 1 X PBS, 50 mM NaF, 0.5% Na deoxycholate (w/v), 0.1% SDS, 1% IGEPAL, 1.5 mM Na3VO4, 1 mM PMSF and complete protease inhibitor (Roche Molecular Biochemicals, IN, USA). The samples were kept on ice and homogenized using a homogenizer. After the homogenization, the sample mixtures were incubated on ice for 10min followed by centrifugation at 10,000 rpm for 10 min at 4°C. The supernatant was collected in a sterile Eppendorf tube and was quantitated with Bio-Rad protein micro assay using BSA as standard (Cat no. 500- 0001). The protein sample of 1ul and the corresponding amount of BSA standard were added to Tris-Hcl solution and then to Bio-Rad dye on a micro titer plate. The plate was then incubated in the spectrophotometer for 30min and the absorbance at 595 nm was recorded. The OD value of the sample and BSA standard were plotted, and the concentration of samples was determined. Based on the concentration, each sample was prepared (5 ug/1 ul) for western blot by adding the sample to 4 X loading dye and heated the mixture at 100°C for 10 min.

Western blotting was done using 10% SDS-PAGE gels. The protein amount of 20 to 80 ug was loaded per lane depending on the target protein and experiments. The SDS PAGE gels were subjected to electrophoresis at 90 V for 90 min with 1 X MOPS running buffer. 10 µl of prestained protein ladder was loaded along with the sample to identify the molecular weight of proteins of interest. The bromophenol blue in the loading buffer was used as the tracking dye. Once the run was complete, the gel was transferred to a PVDF membrane (0.4 um) by wet transfer. The gel, PVDF membrane along with the filter papers and sponges were arranged as a sandwich and placed in the transfer tank with 1 X transfer buffer. The wet transfer was carried out at 100 V for 1 hr or at 30 V for overnight for high molecular weight proteins. On completion of the transfer, the membrane was stained with ponceau stain to check for proper transfer of bands on the membrane. The membrane was blocked with 5% nonfat dry milk in TBST. The primary antibody was added to the 5% nonfat dry milk on the membrane at respective concentration and incubated overnight at 4°C with shaking. The membrane was washed with 5%milk three times for 10 min each and incubated with secondary antibody for 1 hr at room temperature on a shaker. After the incubation, the membranes were washed three times with 5% milk for 10 min each. In the last step, the detection of chemiluminescence was achieved by incubation of the membrane with a substrate such as SuperSignal™ West Femto Maximum Sensitivity Substrate (34094, Thermo Scientific) or SuperSignal™ West Pico PLUS Chemiluminescent Substrate (34577, Thermo Scientific). The substrate was removed, and the membrane was visualized using the Bio-Rad chemiDoc system (Biorad #12003153). Information of the specific antibodies used at 1:1000 dilution:): rabbit anti-GAPDH (ABClonal #A19056), rabbit anti-GR (ABClonal #A2164), rabbit anti-HISTONE H3 (ABClonal #A20822), rabbit anti-FOXC1 (ABClonal #A2924), rabbit anti-ARID5a (Invitrogen MA518292), rabbit anti-Phospho IRS (S307) (ABClonal #AP0371), rabbit anti-IRS (ABClonal #A19245), rabbit anti-AKT (ABClonal #A22533), rabbit anti-Phospho AKT (s473) (ABClonal #AP0098), rabbit anti-GLUT4 (ABClonal #A7637), rabbit anti-INSR (ABClonal #A16900), rabbit anti-FABP4 (ABClonal #A11481), rabbit anti-CD36 (ABClonal #A5792), rabbit anti-HDAC1 (ABClonal #A0238), rabbit anti-SIN3A (ABClonal #A1577), rabbit anti-MYH4 (ABClonal #A15293), rabbit anti-MYH2 (ABClonal #A15292), rabbit anti-MYH7 (ABClonal #A7564), mouse anti-OXOPHOS (Abcam #ab110413). rabbit anti-PPARG (Invitrogen PA3-821A), rabbit anti-SAP30 (Invitrogen PA5-103284. Secondary antibody (diluted 1:3000 dilution): HRP-conjugated donkey anti-rabbit or anti-mouse (#sc-2313 and #sc-2314, Santa Cruz Biotech, Dallas, TX).

Co-immunoprecipitation analysis from the total protein isolated from muscle was assessed for protein complex interaction and difference in interaction among each group of GR ^wt/wt^ and GR ^R24K/R24K^ mice at adult. The Co-immunoprecipitation (Co-IP) protocol includes pulling down the protein complex with an antibody against one member of the complex and coupling the antibody to a magnetic bead, followed by the isolation and elution of the complex and then verification by western blot analysis of each protein complex moieties. The universal magnetic Co-IP kit (Active motif, 54002, Carlsbad, CA, USA) was used and the appropriate antibodies (specified in table 2) and the control IgG of 2 µg were used to pull down the complex. The total protein extract of 800 µg was prepared in a final volume of 500 ul, with the complete Co-IP/Wash buffer and incubated with the specific antibodies and IgG control overnight at 4°C on a rotator. After the incubation, the protein G magnetic beads (Invitrogen # 80105G) were added to the mixture and incubated at room temperature for 1 hr. The magnetic beads were then separated using a magnetic separator and the mixture was discarded. The magnetic beads which hold the corresponding complex were then washed with IP wash buffer three times and the final elution was done by suspending the beads in 50 µl of 2 X loading dye. The beads were headed at 100°C for 5 min. The protein complexes and the beads were then separated using the magnetic stand and loading dye with the proteins were separated into a new tube and the samples were loaded onto a 10% SDS-PAGE gel with 20 µl loaded each lane.

### Nuclear, cytoplasmic and membrane fraction analysis

The separation of nuclear and cytoplasmic protein analysis was performed using NE-PER nuclear and cytoplasmic extraction kit (Invitrogen #78835). Briefly, 100mg of the skeletal muscle was homogenized and 1ml of CERI solution was added and vortexed vigorously on high setting for 15sec. Following incubation on ice for 10 min 55ul of ice cold CERII solution was added, vortexed, incubated for a minute and centrifuged for 5 min at 16,000g. The supernatant containing the cytoplasmic fraction was separated into a new 1.5 Eppendorf tube and then suspended the insoluble pellet with 500ul of ice-cold NER solution. The sample was placed on ice for 40 minutes and vortexed every 10 for 1 sec. Finally, the samples were centrifuged for 10 min at 16,000g and the supernatant containing the nuclear fraction was separated and in a new 1.5 Eppendorf tube and stored at - 80C until use.

The isolation of membrane proteins was achieved by a modified protocol ^66^. Muscle tissue (50 mg) from day 1 ABW and LBW pups were taken and cut into pieces using a sterile blade and transferred to 2 ml Eppendorf tube containing the homogenizing buffer [39 ml Buffer A (121.10 mg Tris-base, 37.22 mg EDTA per 100 ml of dd H2o, at pH 7.4), 13 ml of 20 µm EDTA in buffer A and 312 µl of PMSF], 3 ml of buffer 1 (43.5 g KCl, 13.0 g tetra-sodium pyrophosphate in 500 ml of dd H20). The tissue was homogenized using a homogenizer and the mixture was incubated on ice for 15 min. After the incubation, the samples were centrifuged in an ultracentrifuge at 50,000 rpm for 45 min at 4oC. The pellet was washed in 1 ml of buffer 2 (121.10 mg Tris-base, 37.22 mg EDTA in 100 ml of dd H2O at pH 7.4) and the solution was discarded without disturbing the pellet and the tube was dried with a cotton bud. The pellet was homogenized in 600 µl buffer 2, 200 µl 16% SDS was added and centrifuged at 3000 rpm for 20 min at 20□ C. The supernatant was collected, and the protein concentration was determined by Bio-Rad protein assay as described previously. Once the protein concentration was determined, western blotting analysis was carried out as previously described.

### Protein Immunoprecipitation following LC-MS/ MS analyses

Immunoprecipitation of proteins without the contaminant of antibody heavy and light chain through bead antibody conjugation was carried out using the Pierce Co-IP kit (Invitrogen #26149). Briefly, the antibody of 10-75ug was conjugated with the amino link plus coupling resin using the coupling buffer containing the sodium cyanoborohydride as conjugation reagent was performed in 1.5ml Eppendorf tube in a thermomixer incubated at room temperature for 2 hours. Simultaneously, the protein extracts were pre-cleared with control agarose resins for 1 hour. Then resin was washed with serial solutions of quenching and wash buffers. The eluted pre - cleared protein extracts were added onto the antibody conjugated amino link resin at 4C overnight following which the interaction was washed with was buffer the following day and eluted using 50ul of elution buffer. The eluted protein was run on SDS-PAGE silver stained and western analysis were carried out to confirm the existence of antibody elution following which the samples were submitted to the LC-MS/ MS protein core at UC.

The protein samples were dried by speed vac and resuspended in 35 µl of 1X LB. The samples were then run 1.5cm into an Invitrogen 4-12% B-T gel using MOPS buffer with molecular weight marker lanes in between. The sections were excised, reduced with DTT, alkylated with IAA and digested with trypsin overnight. The resulting peptides were extracted and dried by speed vac. They were then resuspended in 0.1% Formic acid (FA). 500ng-2 ug of each sample was analyzed by nano LC-MS/MS (Orbitrap Eclipse) and was searched against a combined database of a combined contaminants database and the Swissport Mus musculus database using Proteome discoverer version 3.0 with the Sequest HT search algorithm (Thermoscientific).

### Immunostaining

Excised muscle tissues were fixed in 10% formaldehyde (Cat #245-684; Fisher Scientific, Waltham, MA) at room temperature for ∼24 hours, then stored at +4C before processing. Tissue sections of 5–7□µm thickness of was stained with hematoxylin and eosin (H & E; cat #12013B, 1070C; Newcomer Supply, Middleton, WI). CSA quantitation was conducted on >400 myofibers per tissue per mouse. Imaging was performed using an Axio Observer A1 microscope (Zeiss, Oberkochen, Germany), using 10X and 20X (short-range) objectives. Images were acquired through Gryphax software (version 1.0.6.598; Jenoptik, Jena, Germany) and quantitated through ImageJ ^67^. In case of myofiber typing, sections were incubated with primary antibodies BA-F8 (1:10), SC-71 (1:30) and BF-F3 (1:10; all by Developmental Studies Hybridoma Bank, Iowa City, IA) overnight at 4°C. Then, sections were incubated with secondary antibodies AlexaFluor350 anti-IgG2b, AlexaFluor488 anti-IgG1 and AlexaFluor594 anti-IgM (Cat #A21140, A21121, 1010111; Life Technologies, Grand Island, NY). Type 1 fibers stained blue, type 2A stained green, type 2X showed no staining, type 2B stained red. Myofiber types were then quantitated over at least five serial sections and quantitated as % of total counted myofibers.

### Cell culture and gene overexpression analyses

The C2C12 skeletal muscle cell lines, both the wild type and transfected were maintained on DMEM supplemented with 10% fetal bovine serum and 1% Pen strep in 5%CO2 at 37C incubator. When the cells reached 80% confluency, they were transfected using the Lipofectamine 3000 transfection reagent (Invitrogen #L3000015) with plasmid carrying the gene of interest (Foxc1 and Arid5a). After 48 hours of transfection the cells were washed with PBS and RNA and Proteins were extracted as described and assessed for gene of interest overexpression and its associated targets.

### Analyses of body composition and muscle function

Our routine procedures concerning body composition, muscle function, mass and myofiber typing can be found as point-by-point protocols here ^68^.

Forelimb grip strength was monitored using a meter (#1027SM; Columbus Instruments, Columbus, OH) blinded to treatment groups. Animals performed ten pulls with 5 seconds rest on a flat surface between pulls. Grip strength was expressed as force normalized to body weight. Running endurance was tested on a motorized treadmill with electrified resting posts (#1050RM, Columbus Instruments, Columbus, OH) and 10° inclination. Speed was accelerated at 1m/min2 starting at 1m/min and individual test was interrupted when the subject spent >30sec on resting post. Running endurance was analyzed as weight-normalized cumulative work (mW)^69^.

Immediately prior to sacrifice, in situ tetanic force from tibialis anterior muscle was measured using a Whole Mouse Test System (Cat #1300A; Aurora Scientific, Aurora, ON, Canada) with a 1N dual-action lever arm force transducer (300C-LR, Aurora Scientific, Aurora, ON, Canada) in anesthetized animals (0.8 l/min of 1.5% isoflurane in 100% O2). Specifications of tetanic isometric contraction: initial delay, 0.1 sec; frequency, 200Hz; pulse width, 0.5 msec; duration, 0.5 sec; stimulation, 100mA ^70^. Muscle length was adjusted to a fixed baseline of ∼50mN resting tension for all muscles/conditions. Force-frequency curve was measured from 25 Hz to 200 Hz with intervals of 25 Hz, pause 1 minute between tetani. Fatigue analysis was conducted by repeating tetanic contractions every 10 seconds until complete exhaustion of the muscle (50 cycles). Specific force was calculated (N/mm2) for each tetanus frequency as (P0 N)/[(muscle mass mg/1.06 mg/mm3)/Lf mm]. 1.06 mg/mm3 is the mammalian muscle density. Lf=L0*0.6, where 0.6 is the muscle to fiber length ratio in tibialis anterior muscle ^71^. We reported here specific force values in N/cm2 units.

Magnetic resonance imaging (MRI) scans to determine lean mass ratios (% of total body mass) were conducted in non-anesthetized, non-fasted mice at ZT8 using the EchoMRI-100H Whole Body Composition analyzer (EchoMRI, Houston, TX). Mice were weighed immediately prior to MRI scan. Before each measurement session, the system was calibrated using the standard internal calibrator tube (canola oil). Mice were scanned in sample tubes dedicated to mice comprised between 20 g and 40 g body mass. Data were collected through built-in software EchoMRI version 140320. Data were analyzed when hydration ratio > 85 %. Muscle mass was calculated as muscle weight immediately after sacrifice and explant, normalized to whole body weight.

### Respirometry with isolated mitochondria and muscle tissue

Basal tissue OCR values were obtained from basal rates of oxygen consumption of muscle biopsies at the Seahorse XF HS Mini Extracellular Flux Analyzer platform (Agilent, Santa Clara, CA) using previously detailed conditions ^70^. Basal OCR was calculated as baseline value (average of 3 consecutive reads) minus value after rotenone/antimycin addition (average of 3 consecutive reads). Basal OCR values were normalized to total protein content, assayed in each well after the Seahorse through homogenization and Bradford assay. Nutrients: 5mM glucose, 1mM palmitate-BSA (#G7021, #P0500; Millipore-Sigma, St Louis, MO); inhibitors: 0.5mM rotenone + 0.5mM antimycin A (Agilent).

Respiratory control ratio (RCR) values were obtained from isolated mitochondria from muscle tissue. Quadriceps are harvested from the mouse and cut into very fine pieces. The minced tissue is placed in a 15mL conical tube (USA Scientific #188261) and 5mL of MS-EGTA buffer with 1mg Trypsin (Sigma #T1426-50MG) is added to the tube. The tube is quickly vortexed, and the tissue is left submerged in the solution. After 2 minutes, 5mL of MS-EGTA buffer with 0.2% BSA (Goldbio #A-421-250) is added to the tube to stop the trypsin reaction. MS-EGTA buffer: Mannitol-ChemProducts #M0214-45, Sucrose-Millipore #100892, HEPES-Gibco #15630-080, EGTA-RPI #E14100-50.0. The tube is inverted several times to mix then set to rest. Once the tissue has mostly settled to the bottom of the tube, 3mL of buffer is aspirated and the remaining solution and tissue is transferred to a 10mL glass tissue homogenizer (Avantor # 89026-382). Once sufficiently homogenized the solution is transferred back into the 15mL conical tube and spun in the centrifuge at 1,000g for 5 minutes at 4 degrees Celsius. After spinning, the supernatant is transferred to a new 15mL conical tube. The supernatant in the new tube is then centrifuged at 12,000g for 10 minutes at 4 degrees Celsius to pellet the mitochondria. The supernatant is discarded from the pellet and the pellet is then resuspended in 7mL of MS-EGTA buffer and centrifuged again at 12,000g for 10 minutes at 4 degrees Celsius. After spinning, the supernatant is discarded, and the mitochondria are resuspended in 1mL of Seahorse medium (Agilent #103335-100) with supplemented 10µL of 5mM pyruvate (Sigma #P2256-100G) and 10µL of 5mM malate (Cayman Chemical #20765). After protein quantitation using a Bradford assay (Bio-Rad #5000001), 2.5µg mitochondria are dispensed per well in 180µl total volumes and let to equilibrate for 1 hour at 37°C. 20µL of 5mM ADP (Sigma #01905), 50µM Oligomycin (Millipore #495455-10MG), 100µM Carbonyl cyanide-p-trifluoromethoxy phenylhydrazone (TCI #C3463), and 5µM Rotenone (Millipore #557368-1GM)/Antimycin A (Sigma #A674-50MG) are added to drug ports A, B, C, and D respectively to yield final concentrations of 0.5mM, 50µM, 10µM, and 0.5µM. Nutrients: 0.5mM pyruvate, 0.1mM palmitoyl carnitine (#P2256, #61251; Millipore-Sigma, St Louis, MO). At baseline and after each drug injection, samples are read three consecutive times. RCR was calculated as the ratio between state III (OCR after ADP addition) and uncoupled state IV (OCR after oligomycin addition). Seahorse measurements were conducted blinded to treatment groups.

### AAV preparation

Approximately 70-80% confluent HEK293T cells (AAVpro® 293T Cell Line; Takara # 632273 AAVpro® 293T Cell Line; Takara # 632273) in DMEM (SH30022.01, Cytiva Life Sciences) supplemented with 2% Bovine Growth Serum (BGS; Cytiva Life Sciences), and 1.0 mM Sodium Pyruvate were triple transfected with pHelper (Cell Biolabs;340202), pAAV-GOI (Vector Builder; (VB230825-1437xmg; pAAV[Exp]-CMV>{mFoxc1[NM_008592.2]*-3xFLAG:WPRE), VB230825-1437xmg; pAAV[Exp]- CMV>{mArid5a[NM_001290726.1]*-3xFLAG:WPRE)) and pAAV Rep-Cap (1A-Myo; Gift of Molkentin Lab) plasmids using PEI, Linear, MW250,000 (PolySciences, Inc) in 40-T150mm cell culture plates. Eighteen hours after transfection, medium is changed to DMEM supplemented with 1% BGS, 1.0 mM Sodium Pyruvate, and 1X MEM Non-essential Amino Acid Solution (Sigma; M7148). Approximately 96 hours post-transfection, the media and cells were collected and processed separately. Cells were lysed using repeated freeze/thaw cycles at a minimum of five times in 1X Gradient Buffer (0.1 M Tris, 0.5 M NaCl, 0.1 M MgCl2). The cell debris were then treated with Benzonase Endonuclease at 0.65 µl per 5 mL (Sigma-Aldrich #1037731010 (100000 Units)) for at least one hour. The homogenates were cleared from debris by centrifugation. AAVs were precipitated from the cell medium with polyethylene glycol (PEG) 8000 The PEG-precipitated AAV was collected by centrifugation, and the AAV pellet was resuspended in 1X GB. Media and cell AAV’s were combined and AAV’s were purified using an Iodixanol (Opti Prep Density Gradient Medium; Sigma-Aldrich #D1556250) gradient at 15%, 25%, 40% and 60% in 1XGB. The AAV band was removed and purified using Centrifugal Filters (30000 NMWL (30K), 4.0 mL Sample Volume; Millipore-Sigma #UFC803024, and 100000 NMWL (100K), 15.0 mL Sample Volume; Millipore-Sigma # UFC910024) in a2X PBS, 10mM MgCl2 solution.

### Viral titration

Primer’s binding within the AAV-GOI ITR’s CMV region (Forward: GTTCCGCGTTACATAACTTACGG; Reverse: CTGCCAAGTGGGCAGTTTACC) were used to measure the virus titer with quantitative polymerase chain reaction (qPCR). Before releasing the viral DNA from the particles, all extra-viral DNA was removed by digestion with DNase I. Then, the viral DNA was released by Proteinase K digestion.

### In vivo viral injection

The viral load of MyoAAV’s corresponding to 10˄12 per construct per mouse were administered retro-orbitally in anesthetized mice. Muscles were then excised after 2 weeks for immediate sufficiency proofs, or after 12 weeks in combination with high-fat diet for sufficiency proofs in the presence of metabolic stress.

### UK Biobank and All of Us analyses

Our analyses were conducted under the UKB application number 65846 and All of Us workspace number aou-rw-0fb52975. We constructed a rs6190 genotype-stratified cohort, excluding participants if they withdrew consent. All available values for the tested parameters were collected per genotype group. For UK Biobank, UDI and related parameters: Age: 21001-0.0; BMI: 21001-0.0; Glycemia (mM): 30740-0.0; Triglycerides (mM): 30870-0.0; lean mass (kg): 23280-0.0; hand grip strength (kg): 46-0.0, 47-0.0. Regression analyses were performed using second generation of PLINK ^72^. Before analyses, a series of standard QC measures were applied including sample call rates, sample relatedness, and sex inconsistency as well as marker quality (i.e., marker call rate, minor allele frequency (MAF), and Hardy-Weinberg equilibrium (HWE). Analyses were limited to participants with call rates >□98%, SNPs with call rates >□99%, and SNPs with MAF >□1% and HWE p□>□0.0001. For independent association confirmation studies, multiple linear regression analysis was carried out using R 4.3.2 (R Core Team, 2023) to explore the association of target metabolic parameters versus rs6190 genotype and correcting for 10 principal components, triglyceridemia, sex ratio and age.

### Statistics

Unless differently noted, statistical analyses were performed using Prism software v8.4.1 (GraphPad, La Jolla, CA). The Pearson-D’Agostino normality test was used to assess data distribution normality. When comparing the two groups, a two-tailed Student’s t-test with Welch’s correction (unequal variances) was used. When comparing three groups of data from one variable, one-way ANOVA with Sidak multi-comparison was used. When comparing data groups for more than one related variable, two-way ANOVA was used. For ANOVA and t-test analyses, a P value less than 0.05 was considered significant. When the number of data points was less than 10, data were presented as single values (dot plots, histograms). Tukey distribution bars were used to emphasize data range distribution in analyses pooling larger data points sets per group (typically > 10 data points). Analyses pooling data points over time were presented as line plots connecting medians of box plots showing distribution of all data per time points. Randomization and blinding practices are followed for all experiments. All the data from all animal cohorts and cell clone replicates is reported, whether outlier or not.

## Data availability

RNA-seq and ChIP-seq datasets reported here are available on GEO as entries GSE262234 and GSE262235.

---

**Supplementary Figure 1.**
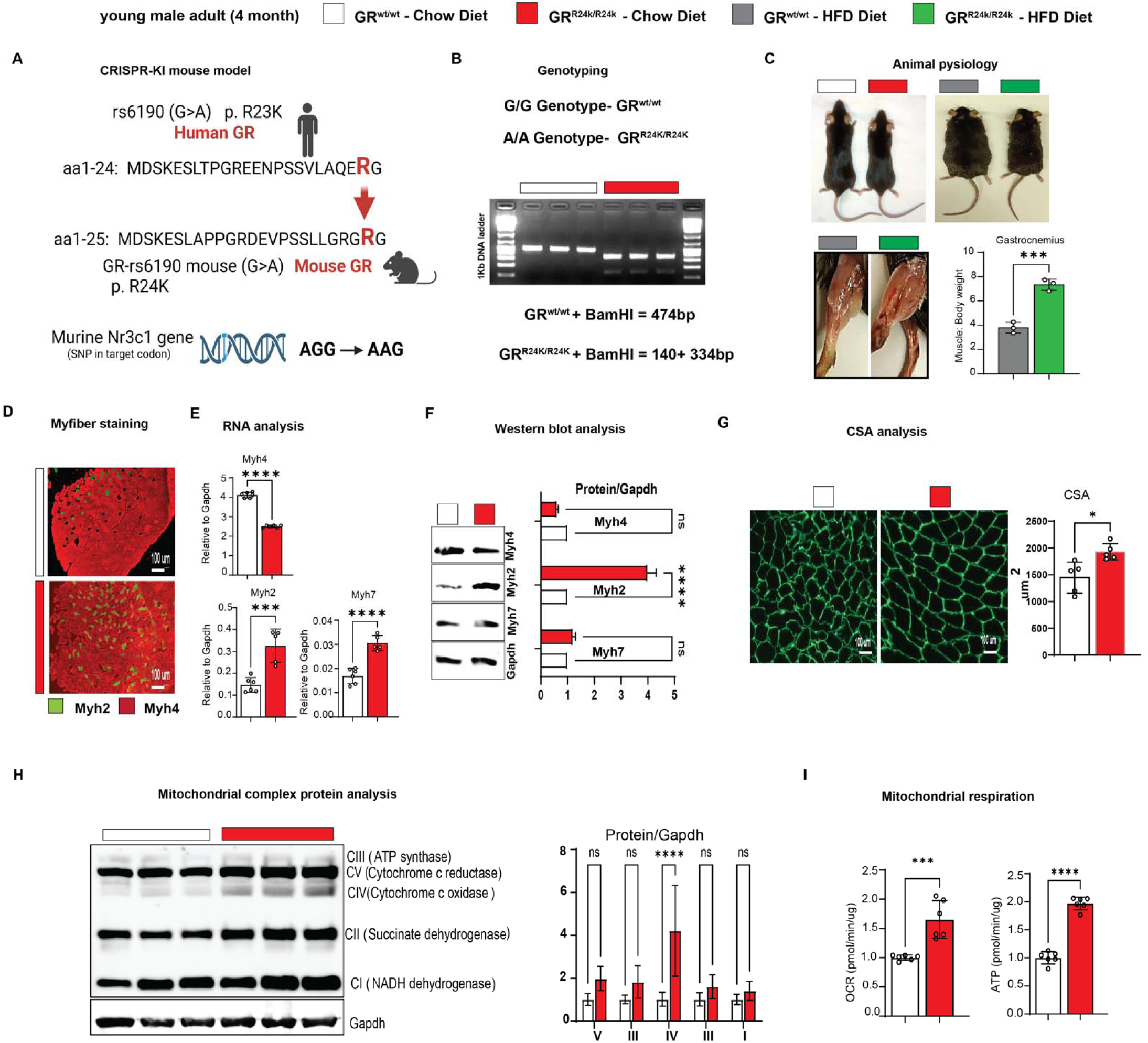
Additional data regarding R24K effects on muscle metabolism. Related to. Figure 1 **– (A-B)** Diagram and genotyping PCR for R24K genocopying strategy in mice of human R23K variant. **(C)** Muscle/body mass was increased by the R24K genotype in obese mice. **(D-F)** R24K muscles showed increased expression of Myh2 (type 2A) and Myh7 (type 1) myosins, with decreased Myh4 (type 2B) expression. **(G)** Cross-sectional area (CSA) analysis confirmed increased muscle mass. **(H)** Mitochondrial complex protein levels in muscle showed non-significant upward trends, with complex IV bing significantly upregulated by the R24K homozygosity. **(I)** Glucose-fueled respiration was increased in muscle tissue of R24K vs WT mice. n=3-6♂/group; diet exposures for 12 weeks from 4mo to 7mo; Welch’s t-tests: ns, non significant; *, P<0.05; **, P<0.01; ***, P<0.001; ****, P<0.0001.

**Supplementary Figure 2.**
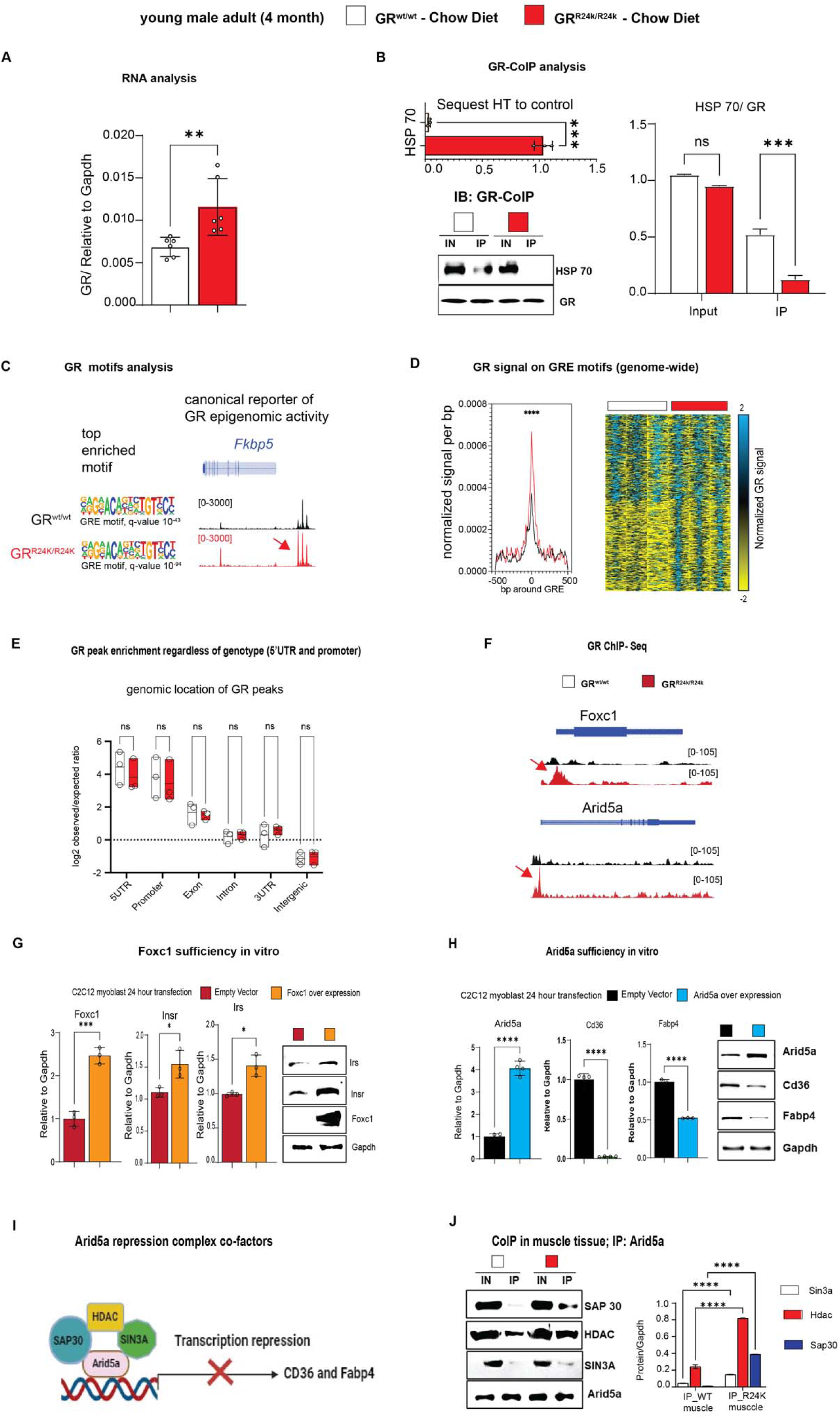
Additional analyses related to the mutant GR in muscle –. **(A)** Validation of lower Hsp70 binding by the mutant vs WT GR through CoIP. **(B)** qPCR analysis of *Nr3c1* (GR gene) expression in muscle. **(C)** Validation of ChIP-seq datasets through unbiased motif analysis and visualization of canonical GR peaks in the *Fkbp5* promoter region. **(D-E)** Mutant GR epigenomic activity was increased on the GR-binding elements (GREs) genome-wide, despite no significant shifts in overall genomic location (GR peaks enriched for both genotypes in the promoter-5’UTR regions). **(F)** Mutant vs WT GR peaks on Foxc1 and Arid5a proximal promoters. **(G-H)** Validation of Foxc1 and Arid5a sufficiency for target gene programs in vitro in C2C12 myoblasts. **(I-J)** Diagram and CoIP for Arid5a repression complex partners in R24k vs WT muscle. n=3- 4♂/group; Welch’s t-tests and 2w ANOVA + Sidak (J): ns, non significant; *, P<0.05; **, P<0.01; ***, P<0.001; ****, P<0.0001.

